# What is the influence of knee joint movement on the maximal average mechanical power output of human quadriceps femoris muscle?

**DOI:** 10.1101/2025.11.23.689993

**Authors:** Edwin D.H.M. Reuvers, A.J. ‘Knoek’ van Soest, Dinant A. Kistemaker

**Affiliations:** Department of Human Movement Sciences, Faculty of Behavioural and Movement Sciences, Vrije Universiteit Amsterdam, Amsterdam Movement Sciences, The Netherlands

**Author notes:** Corresponding author: Edwin D.H.M. Reuvers. Access the code and full analysis notebook at https://edwinreuvers.github.io/publications/hq-ampo.

## Abstract

Performance during motor activities such as wheelchair riding, cycling, rowing and speed skating, depends critically on the average mechanical power output (AMPO) produced by the muscles. To maximise short-duration performance, limb movements should allow muscles to deliver maximal AMPO. However, it is unclear which movement maximises AMPO of human muscle. In this study, we employed a Hill-type muscle-tendon-complex (MTC) model to predict the maximally attainable AMPO of human *m. quadriceps femoris* for various imposed periodic knee joint movements. Based on these predictions, we selected one set of conditions predicted to yield identical maximally attainable AMPO despite substantial variations in knee joint movements and another set of conditions predicted to yield substantial variations in maximally attainable AMPO. In the experiment, periodic knee joint movements were fully imposed by a knee dynamometer. Participants were instructed to maximise AMPO and, to this end, received visual feedback on their cumulative mechanical work throughout each cycle. Experimental data closely matched predictions derived from the Hill-type MTC model, confirming the validity of the model. Model predictions showed a substantial influence of knee joint movement on the maximally attainable AMPO. Specifically, predictions revealed a strong interaction between cycle frequency and knee joint excursion: increasing one necessitates a decrease in the other to maximise AMPO. Even more interestingly, *m. quadriceps femoris* should spend about 80% of the cycle duration while shortening, independent of cycle frequency and/or knee joint excursion.

## 1 Introduction

Basic motor activities such as walking, running, swimming, and activities involving man-made appliances like wheelchair riding, cycling, rowing, and speed skating are characterised by repetitive limb movements that result in periodic stretch-shortening cycles (SSCs) of muscles. In each SSC, muscles alternate between concentric and eccentric phases: they deliver positive mechanical power when generating force while shortening, and they deliver negative mechanical power when generating force while lengthening. In many motor activities, performance largely depends on the average mechanical power output (AMPO) over a full periodic cycle of all muscles involved. Maximising performance therefore requires the muscles to maximise average positive mechanical power during shortening and to minimise average negative mechanical power during lengthening. This raises the question: what periodic muscle length/velocity over time should be imposed in order to maximise AMPO?

A muscle consists of contractile fibres (the contractile element; CE) and connective tissue, both parallel to the fibres (the parallel elastic element; PEE) and in series with the fibres (the series elastic element; SEE). These three elements together form the muscle-tendon complex (MTC). Because connective tissue behaves predominantly elastically, AMPO of the MTC is determined almost exclusively by AMPO of the CE. However, CE length depends on MTC length and deformation of the elastic elements, which in turn depends on MTC force. Consequently, and ultimately, MTC AMPO depends not only on stimulation over time but also on MTC length over time.

During limb movement, MTC length varies with the joint(s) it spans. Understanding how periodic MTC length changes affect AMPO thus provides insight into how joints should move to maximise AMPO. These insights are not only interesting from a fundamental scientific perspective, but also from an applied sciences perspective, as they can inform the (re-)design of equipment. Such a design approach, that capitalises on a thorough understanding of the human musculoskeletal system has proven effective, as demonstrated by the Klapskate for speed skating (Ingen Schenau et al., 1996) and the SuperSeat for rowing (Soest and Hofmijster, 2009). Yet, such design approaches remain rare due to limited understanding of how changes in MTC length over time affect AMPO, or other measures of performance in human muscle.

Much of the existing knowledge of the influence of CE/MTC length over time on AMPO is inferred from experiments on isolated animal muscle. In these experiments, CE or MTC length and muscle stimulation are imposed and resulting AMPO is measured. This method is often referred to as the ‘workloop technique’ (Josephson, 1985). These experiments have often been performed for sinusoidal SSCs, showing that the effects of cycle frequency and CE/MTC length excursion interact: at lower cycle frequencies, length excursions should be larger in order to maximise AMPO. However, sinusoidal SSCs by definition result in equal shortening and lengthening durations, leaving unanswered how unequal shortening and lengthening duration affects AMPO.

Askew & Marsh (1997; 1998) were the first, and until very recently (see Reuvers et al., 2025), the only researchers to perform experiments on the influence of unequal shortening and lengthening duration on the maximally attainable AMPO of muscle during short-duration, all-out (sprint) conditions. In their study, Askew & Marsh (1997; 1998) imposed SSCs with near-constant shortening and lengthening velocities and parametrised them in terms of three variables: cycle frequency, FTS (fraction of the cycle time spent shortening) and length excursion (see Figure 1). Askew & Marsh (1997; 1998) examined three distinct FTS values: 0.25, 0.50 and 0.75. They found that AMPO was consistently highest for FTS = 0.75 at all tested cycle frequencies at the optimal length excursion. In a recent study (Reuvers et al., 2025), we further examined the effect of SSC parameters on the maximally attainable AMPO by combining experiments on rat m. gastrocnemius medialis with predictions using a Hill-type MTC model. Our results showed a near-perfect correlation between model-based predictions and experimental results (r^2^ > 0.98). The effect of cycle frequency, FTS and length excursion on the maximally attainable AMPO was then further examined using the Hill-type MTC model for a broad range of SSC parameters. These model-based predictions showed that the optimal FTS ranged 0.84-0.88 across a wide range of cycle frequencies and length excursions.

**Figure 1:**
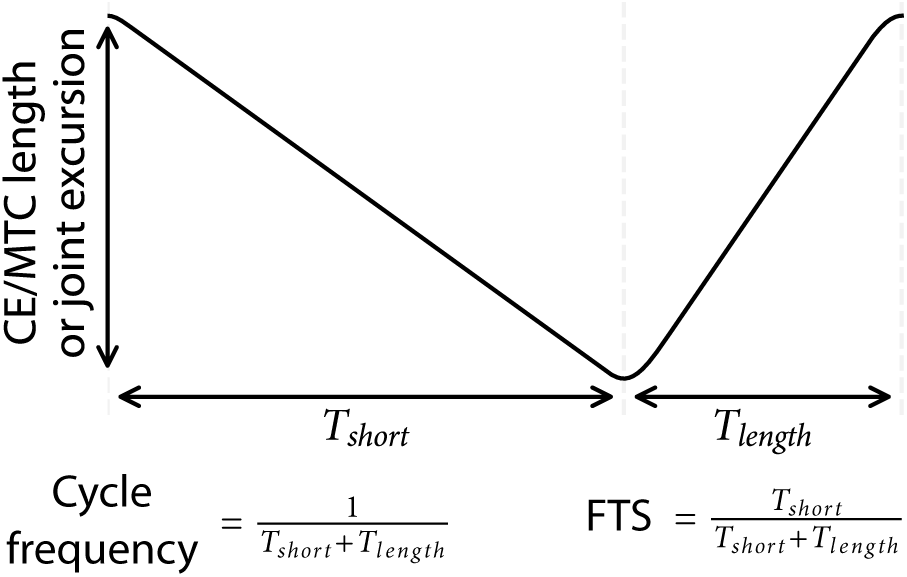
Parameterisation of stretch-shortening cycles. *T_short_* and *T_length_* denote the shortening and lengthening durations. FTS denotes the fraction of the cycle time spent shortening. In the example shown, the shortening duration is 65% of the cycle duration (i.e. FTS = 0.65).

Although animal studies have provided valuable insights, it remains unknown how these findings translate to human muscle. For example, is the optimal FTS for human muscle also in the order of 0.85? Although similar experiments can, in principle, be designed for human muscle, inherent differences exist between experiments involving human participants and experiments on isolated animal muscles. First, MTC properties can be accurately estimated in animal experiments based on dedicated experiments (see Reuvers and Kistemaker, 2025) that are challenging in human experiments (i.e. ‘quick-release’ experiments). Second, CE/MTC length is under full control in animal experiments, whereas in human movement it can only be imposed at the joint level. As a result, the exact CE/MTC length over time is uncertain in human experiments due to inter-individual variation in, for example, tendon compliance and moment arms. Third, stimulation is also under full control of the experimenter in animal experiments, but not in human experiments. Despite these differences, it is feasible to design human experiments in which imposed joint movements are systematically varied, with participants aiming to maximise AMPO. Such an approach allows investigation of the effects of SSC parameters – cycle frequency, FTS, and joint excursion – on maximally attainable AMPO. Surprisingly, to our knowledge such studies have not yet been conducted for human muscle – not even for sinusoidal SSCs. Consequently, the effect of SSC parameters on the maximally attainable AMPO in human muscle is currently unknown.

In this study, we examine the influence of SSC parameters on the maximally attainable AMPO of human *m. quadriceps femoris* during short-duration, all-out (sprint) conditions. Under such conditions, performance is determined by the mechanical properties of the MTC. Hill-type MTC models have been shown to accurately predict mechanical behaviour under these conditions (e.g. Ingen Schenau et al., 1988; Lemaire et al., 2016; Reuvers et al., 2025; Sandercock and Heckman, 1997; Wakeling and Johnston, 1999; Zandwijk et al., 1996). We therefore combined model predictions with *in vivo* knee dynamometer experiments. The model was used to predict the maximally attainable AMPO across a wide range of imposed knee joint movements, from which nine insightful experimental conditions were selected. In the experiment, knee joint movements were fully imposed by a knee dynamometer, while participants aimed to maximise AMPO. Measured AMPO matched the model predictions reasonably well, confirming that the Hill-type MTC model reliably captures the influence of SSC parameters on the maximally attainable AMPO. This justified using the model to systematically examine the overall influence of imposed knee joint movement on the maximally attainable AMPO of human *m. quadriceps femoris*.

## 2 Methods

In this study, we examined the influence of SSC parameters – cycle frequency, FTS, and joint excursion – on the maximally attainable AMPO of human *m. quadriceps femoris*. Experiments were combined with simulations and optimisations using a Hill-type MTC model. Based on model predictions, we selected one set of conditions expected to yield similar AMPO despite substantial variations in knee joint movements, and another set expected to yield large differences in AMPO. In the experiment, knee joint movements were fully imposed by a dynamometer. Participants were instructed to maximise AMPO and, for this purpose, received visual feedback on cumulative mechanical work throughout each cycle. Measured AMPO was compared with model predictions. After confirming that measurements closely matched predictions, the model was used to investigate the overall influence of SSC parameters on the maximally attainable AMPO of human *m. quadriceps femoris*.

### 2.1 Experiment

#### 2.1.1 Participants and ethical approval

Approval of the local ethics committee was obtained before the start of the *in vivo* experiment (VCWE-2016-135). Four male and two female healthy participants (22.8 ± 2.2 years, mean ± SD), who all signed informed consent, participated in the experiment.

#### 2.1.2 Measurement setup

A kinematically driven knee dynamometer (custom made, Vrije Universiteit, Amsterdam, the Netherlands, see Ruiter et al., 2006) was used to fully impose periodic knee joint angle over time. This dynamometer was capable of imposing large knee joint accelerations and was sufficiently stiff such that participant behaviour (i.e. knee joint moment changes) did not influence the resulting knee joint movement (see Results). During the experiment, participants sat in the dynamometer with the trunk, hip, and thigh firmly strapped. The foot was secured to the dynamometer footplate to fix the ankle joint. The fixed joint and segment angles were as follows: the hip angle (between trunk and thigh, 0 rad = fully extended) was about 1.2 rad, the ankle angle (between shank and foot, 0 rad = fully extended) was about 1.2 rad and the thigh angle (relative to the horizontal) was about 0.2 rad (see Figure 2). The knee joint angle was defined such that 0 rad corresponded to a fully extended knee, while knee joint angle increased with flexion (see Figure 2). Participants performed the experiment with their preferred leg. The lower leg was attached to the sliding carriage of the dynamometer just proximal to the lateral malleolus (see Figure 2). To align the knee joint with the rotation axis of the dynamometer, the lateral epicondyle was initially used as a proxy for the knee joint axis while the participant generated a substantial knee extension moment. Next, the participant actively extended and flexed their knee over the experimental joint angle range. During this movement, the travel of the sliding carriage was visually inspected, and the knee joint alignment was adjusted to minimise carriage displacement. Net knee joint moment and knee joint angle were measured with a torque sensor (custom made, Vrije Universiteit, Amsterdam, the Netherlands) and an angle sensor (custom made, Vrije Universiteit, Amsterdam, the Netherlands) at a sample rate of 1000 Hz.

**Figure 2:**
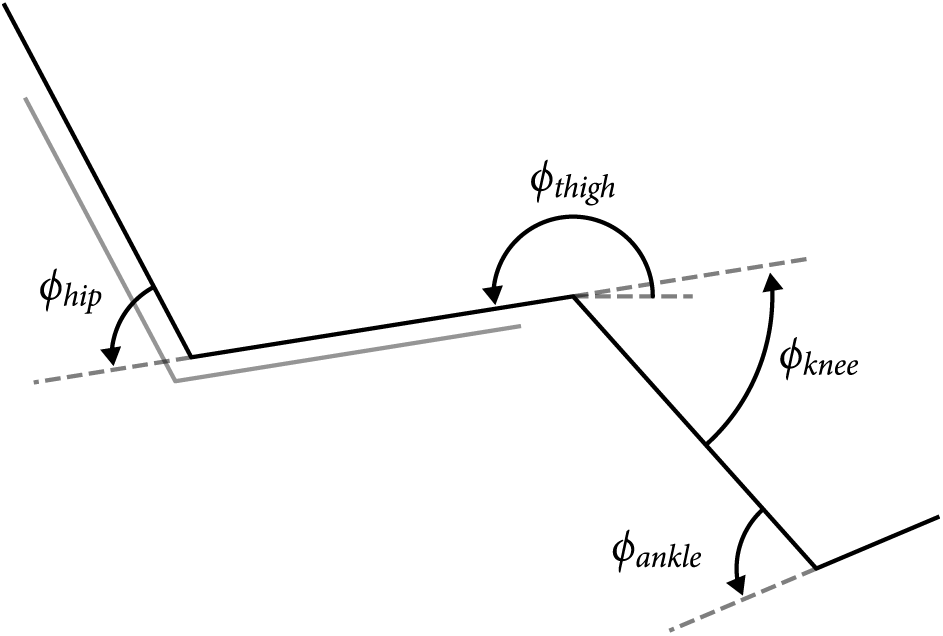
The experimental setup.

Surface EMG of the *m. vastus medialis*, taken as the reference muscle for *m. quadriceps femoris*, was measured during the AMPO measurements. In addition, EMG of *m. biceps femoris*, taken as the reference muscle for the hamstring muscle group, was measured to test for undesired co-contraction. Bipolar EMG electrodes (Ambu, Ballerup, Denmark) were placed at an interelectrode distance of 20mm, at 4/5 of the line originating from the anterior spina iliaca superior to the joint space in front of the anterior border of the medial ligament and at 1/2 of the line originating from the ischial tuberosity to the lateral epicondyle of the tibia for *m. vastus medialis* and *m. biceps femoris* respectively (SENIAM guidelines, Hermens et al., 2000). EMG recordings were amplified with a pre-amplifier (9200 EMG, Goodvaerts Instruments, Kortenhoef, the Netherlands) and collected at a sample rate of 4000 Hz.

#### 2.1.3 Isometric measurements

The maximally attainable AMPO of participants depends on their maximal isometric force. Consequently, for a meaningful comparison between the magnitude of measured AMPO and predicted maximally attainable AMPO, as well as for meaningful comparisons between participants, we needed an accurate estimate of each participant’s maximal isometric force. To obtain this estimate, we measured the isometric knee joint moment-angle relationship for each participant individually. Maximal isometric net knee joint moment was determined at nine different knee joint angles equidistantly distributed over a range of knee joint angles between 0.2 rad and 1.8 rad. The order of these measurements was assigned pseudo-randomly such that they were equally distributed across participants. Participants were instructed to maximise their net knee joint moment with their *m. quadriceps femoris* at every knee joint angle for a period of about five seconds. Participants received real-time visual feedback about their net knee joint moment on a monitor which was placed in front of them and were verbally encouraged by the experimenter. There was a 3-minute rest between each isometric measurement. At the end of the experiment, the isometric measurements for the first two knee joint angles were repeated to assess fatigue.

#### 2.1.4 AMPO measurements

Experimental conditions for the AMPO measurements were selected based on simulations using a Hill-type MTC model. Due to limits on knee angular acceleration, only a subset of combinations of cycle frequency, FTS and knee joint excursion were experimentally feasible (for details, see Section A1). Nine experimental conditions that differed in cycle frequency and FTS at a knee joint excursion of 1.6 rad were used in the experiment (see Figure 3). One subset of five experimental conditions was predicted to yield identical maximally attainable AMPO despite substantial variations in cycle frequency and FTS (0.35A-E; Figure 3). A second subset of five experimental conditions was predicted to yield substantial differences in maximally attainable AMPO due to variations in cycle frequency and FTS (0.20, 0.25, 0.30, 0.35A-E and 0.40; Figure 3). One experimental condition was shared between both subsets, resulting in nine unique experimental conditions.

**Figure 3:**
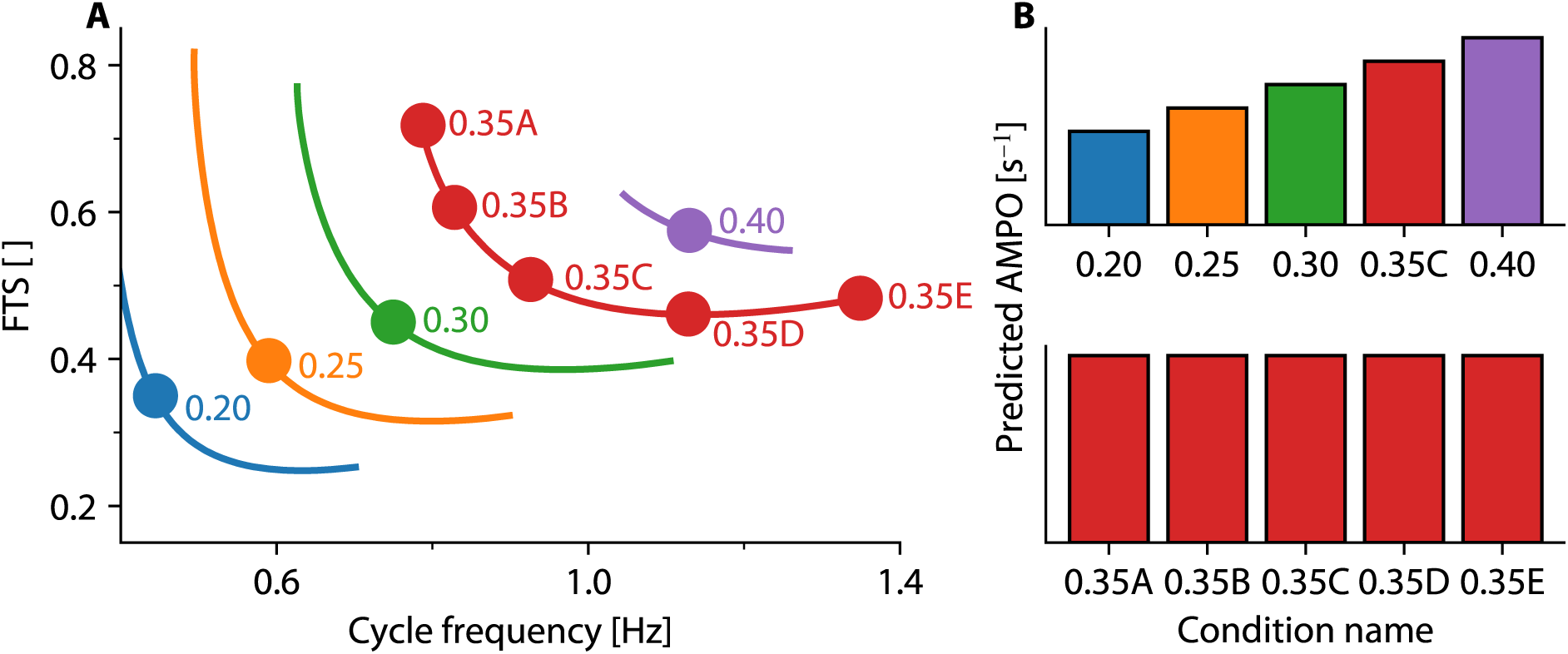
Predicted maximally attainable AMPO and experimental conditions. A) Contour lines depict model-based predictions of maximally attainable AMPO as a function of cycle frequency and FTS for a knee joint excursion of 1.6 rad. Contour lines are only shown for experimentally feasible conditions (due to limits set on knee joint acceleration; see Section A1). Dots indicate the nine conditions selected to be evaluated experimentally. B) Barplots of the same conditions: top, subset predicted to differ in maximally attainable normalised AMPO (i.e. 0.20, 0.25, 0.30, 0.35C and 0.40); bottom, subset predicted to yield identical maximally attainable normalised AMPO (0.35A-E).

In each experimental condition, 15 consecutive cycles of the respective knee joint movement were imposed; the participants were instructed to remain fully relaxed during cycles 1-5 and cycles 11-15 and to maximise AMPO with their *m. quadriceps femoris* during cycles 6-10. Throughout all cycles, participants were instructed to relax their hamstring muscle group to prevent it from producing negative and positive mechanical work during knee extension and flexion.

The periodic knee joint angle over time was fully imposed by the knee dynamometer and therefore periodic lengthening and shortening of the MTC was fully imposed. Consequently, participants could only influence measured AMPO through their muscle activation. For the predictions derived with the Hill-type MTC model, muscle activation was optimised for AMPO (for details, see Section 2.2.2). Thus, for a meaningful comparison between measured AMPO and the predicted maximally attainable AMPO, it was crucial that participants chose a (near-)optimal muscle activation of their *m. quadriceps femoris*. Therefore, each participant practised the nine conditions twice during both practice days (day 1 and 2) of the experiment. Additionally, all conditions were performed once at the end of day 3, at which day the isometric measurements took place. Thus, in total, participants practised each of the nine conditions five times, equalling 25 active cycles. During both practice and the actual AMPO measurements, participants received real-time visual feedback about their cumulative work throughout each cycle on a monitor and were verbally encouraged by the experimenter to maximise it.

During the AMPO measurements day (day 4), EMG measurements of m. *m. quadriceps femoris* and the hamstring muscle group were taken. At the start and at the end of the AMPO measurement, participants performed isometric contractions at a knee joint angle of 1.0 rad: twice for *m. quadriceps femoris* and twice for the hamstring muscle group in order to record EMG signals during maximal voluntary contractions (*EMG^MVC^*) of both muscles, which were later used to normalise EMG activity during the AMPO measurements (see Section 2.3). Before the actual measurement, the participants again practised with the protocol by performing the four most extreme conditions of the experiment (i.e. 0.20C, 0.35A, 0.35E and 0.40C). To prevent fatigue during the actual measurements, participants were instructed to perform these conditions sub-maximally. After this, participants performed the nine different conditions, from which the first two were repeated at the end to check for muscle fatigue. As with the isometric measurements, the order of the AMPO conditions was assigned pseudo-randomly such that they were equally distributed across participants. There was a 3-minute rest between each condition.

### 2.2 Musculoskeletal model

#### 2.2.1 Description

A planar model consisting of an upper and lower leg connected by a frictionless hinge joint was used to relate knee joint angle to muscle-tendon-complex (MTC) length and to relate MTC force to knee joint moment (for details, see Section A2). The musculoskeletal model did not contain any knee flexors, as participants were instructed to maximise AMPO using their knee extensors only (see Section 2.1.2). A single Hill-type MTC model was used to represent the mechanical behaviour of all knee extensors in the human leg. The formulation of the Hill-type muscle-tendon complex model has been previously described in full detail (Reuvers and Kistemaker, 2025; Soest and Bobbert, 1993). Briefly, the Hill-type MTC model included a contractile element (CE) with a parallel elastic element (PEE) in parallel and a serial elastic element (SEE) in series with the CE. PEE and SEE were modelled as purely passive elastic structures, with their force non-linearly dependent on their length. CE force depended on CE length, CE velocity and active state. Active state, defined as the relative *Ca*^2+^ concentration bound to troponin C (Ebashi and Endo, 1968), depended on free *Ca*^2+^ concentration and CE length (Hatze, 1981, pp 31-32; Kistemaker et al., 2005). The free *Ca*^2+^ concentration in turn, depended on neural drive (*STIM*) by first-order dynamics. *STIM* and the imposed knee joint angle over time were the model’s independent inputs.

#### 2.2.2 Predictions of maximally attainable AMPO

To derive predictions of the maximally attainable AMPO, we used the musculoskeletal model with parameter values obtained from literature (see Table A1). As such, we derived generic predictions without tuning those to the muscle parameters of the participants in our experiment. Using the musculoskeletal model, we imposed periodic knee joint movements in which knee joint angle was centred around 1.0 rad. We set the knee angular velocities to constant values that could differ between flexion and extension. We examined a large number of knee joint movements over a range of cycle frequencies (0.4-3 Hz, in 0.2 Hz steps), FTS values (0.10-0.90, in 0.05 steps) and knee joint excursions (0.8-2.0 rad, in 0.1 rad steps). Muscle activation onset time (i.e. the time instance at which *STIM* switches from 0 to its maximal value) was chosen at the start of MTC shortening for each imposed knee joint movement. Muscle activation duration (i.e. the time period during which *STIM* remains at its maximal value before returning to 0) was optimised for AMPO using a Nelder-Mead Simplex method (Lagarias et al., 1998). To ensure periodic behaviour, simulations were run for at least five cycles and until the difference in MTC force at the beginning and end of the cycle was less than 0.1% *F^max^_CE_*. AMPO was calculated for the last cycle.

### 2.3 Data analysis

The isometric measurements were used to estimate the maximal isometric force (*F^max^_CE_*) for each participant individually. The maximal net knee joint moment was determined for every measured knee joint angle and was defined as the highest 500 ms average after correcting for gravitational effects. The moment arm was assumed to be equal for all participants (i.e. moment arm = 4.2 cm, see ??). The isometric knee joint moment-angle relationship of the musculoskeletal model depends on three parameters: *F^max^_CE_*, *L^opt^_CE_* (CE optimum length), and *L*^0^_SEE_ (SEE slack length). These parameters were estimated individually for each participant by minimising the sum of squared differences between the measured isometric net knee joint moment and those predicted by the musculoskeletal model. This optimisation was performed using a Nelder-Mead Simplex method (Lagarias et al., 1998).

EMG signals measured during the AMPO measurements were bandpass filtered using a bidirectional first order Butterworth filter with cut-off frequencies of 20 and 500 Hz (Butterworth, 1930). The EMG envelope was determined as the absolute value of the Hilbert transformed EMG signal (Hilbert, 1904). *EMG^MVC^* values of *m. vastus medialis* and *m. biceps femoris* were defined as the maximal average value of the EMG envelope over 500 ms during the isometric contractions at a knee joint angle of 1.0 rad. Measured EMG signals of *m. vastus medialis* and *m. biceps femoris* during the AMPO measurements were smoothed by a moving average filter with a window of 100 ms and were normalised by their *EMG^MVC^* value. The first and last 0.1 seconds of knee extension were not taken into account for the calculation of the average EMG values, as EMG values increased and decreased during these time intervals. Therefore, average EMG values reported were calculated between a knee joint angle of 0.45 and 1.55 rad.

The knee joint moment measured by the dynamometer during the AMPO measurements represented the combined contributions of gravity, inertia (from both the participant’s leg+foot and that of the dynamometer), passive structures, and active muscle force. To estimate the contribution of gravity, inertia and passive structures, participants were instructed to remain fully relaxed during the first and last five cycles of each experimental condition. The knee joint moment recorded during these passive cycles was assumed to reflect only the net non-muscular contribution. To isolate the muscular contribution, the average knee joint moment from passive cycles 2, 3, 4, 12, 13, and 14 was calculated and subtracted from the measured knee joint moment. The resulting moment, attributed to the moment resulting from muscle force only, was called the ‘muscular’ knee joint moment (*M_mus_*). AMPO per individual cycle was calculated by integrating *M_mus_* over knee joint angle using a trapezoid method, and dividing the result by the cycle duration.

Measured AMPO and predicted maximally attainable AMPO were normalised in two different ways. First, we assessed whether the magnitude of measured AMPO could be adequately predicted. The predicted maximally attainable AMPO scales linearly with maximal isometric force (*F^max^_CE_*) and also depends on CE optimum length (*L^opt^_CE_*). The sensitivity for *L^opt^_CE_* was, however, low: a 10% reduction in *L^opt^_CE_* resulted in only a 3.7 ± 0.3% decrease in AMPO across all experimental conditions. We used the product of the generic model parameter value of *L^opt^_CE_* (see Table A1) and each participant’s individually estimated *F^max^_CE_* to normalise measured AMPO. The predicted maximally attainable AMPO was normalised to the generic model parameter values of *L^opt^_CE_* and *F^max^_CE_*. Second, we evaluated how well differences in AMPO between experimental conditions were predicted. For each subject, the mean (*µ*) and standard deviation (*σ*) of measured AMPO across all experimental conditions were calculated, and individual condition values were transformed into z-scores for each condition (denoted by the subscript *i*):

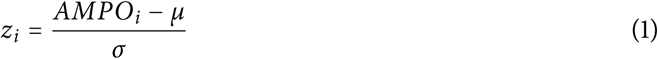

This procedure removed the influence of all potential errors affecting the magnitude of measured AMPO and isolates the relative changes of AMPO. Model-predicted maximally attainable AMPO values were similarly transformed into z-scores, allowing a direct comparison between the measured and predicted z-scores. Importantly, regardless of the normalisation method, whether by the product of *L^opt^_CE_* and *F^max^_CE_* or by z-scores, measured AMPO was assessed relative to the maximally attainable AMPO calculated from the generic model parameters. Thus, normalised model predictions were based on generic model parameters obtained from literature.

## 3 Results

### 3.1 AMPO-scaling factors were successfully obtained

All six participants were able to execute the isometric measurements at all imposed knee joint angles. To test for fatigue, the first two knee joint angles were repeated at the end of the isometric measurements. The maximal knee joint moment was 6.6 ± 5.9% lower for the last two measurements compared to the first two measurements, averaged over all participants and the two repeated measurements. The normalised root mean square difference between measured and predicted isometric net knee joint moments was 5.1 ± 2.4% on average. Variance in the estimated values of *L^opt^_CE_* and *L*^0^_SEE_ between participants was low (Table 1). A representative example of the measured and modelled maximal isometric knee joint moment as a function of knee joint angle is depicted in Figure 4.

**Figure 4:**
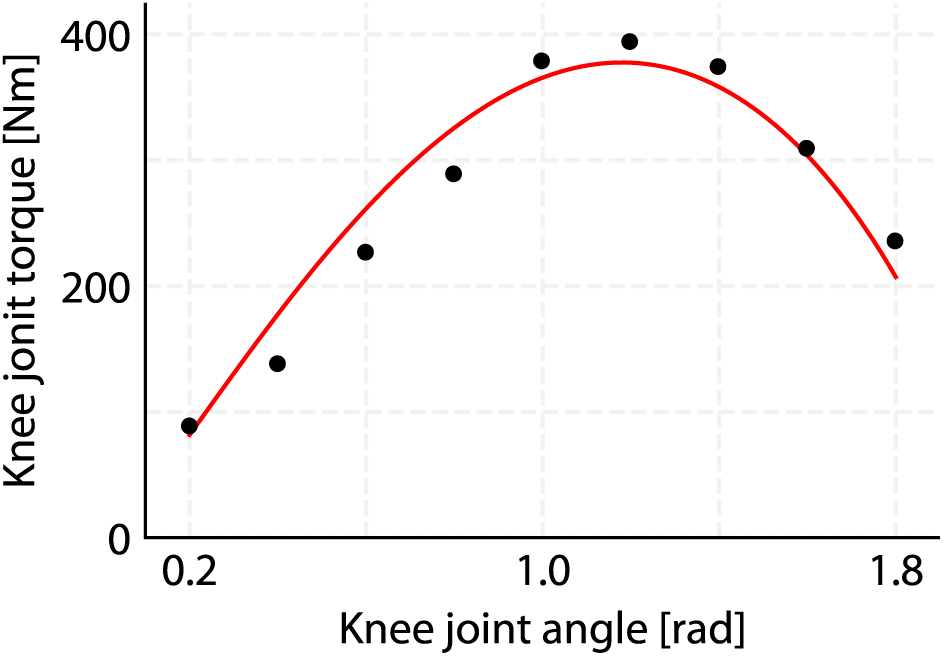
Representative example of maximal isometric knee joint moment as a function of knee joint angle, shown for participant 5. Maximal knee joint moment was measured (black dots) and used to estimate *L*^0^_SEE_, *L^opt^_CE_* and *F^max^_CE_*. The resulting maximal isometric knee joint moment of the musculoskeletal model is depicted by the red line.

**Table 1:** Estimated parameter values for each individual participant.

| Participant | $F_{CE}^{max}$ [N] | $L_{CE}^{opt}$ [cm] | $L_{SEE}^0$ [cm] | RMSD [%] |
| --- | --- | --- | --- | --- |
| 1 | 4500 | 8.5 | 17 | 7.6 |
| 2 | 5200 | 9.7 | 15 | 7.1 |
| 3 | 5700 | 8.6 | 17 | 3.6 |
| 4 | 6800 | 8.7 | 16 | 3.6 |
| 5 | 9100 | 7.5 | 18 | 6.8 |
| 6 | 8500 | 8.9 | 16 | 1.7 |
| AVG±SD | 6600 ± 1900 | 8.6 ± 0.7 | 17 ± 1.1 | 5.1 ± 2.4 |

$F_{CE}^{max}$ , $L_{CE}^{opt}$ and $L_{SEE}^0$ were estimated from the isometric measurements for each participant individually. The root mean square difference (RMSD) was normalised by the individual maximal predicted knee joint moment, which is the product of $F_{CE}^{max}$ and the moment arm of m. quadriceps femoris.

### 3.2 Knee joint movements were successfully imposed

For a meaningful comparison between the measured and predicted maximally attainable AMPO, it was crucial that the knee dynamometer adequately imposed the desired knee joint movement of the experimental conditions. This to ensure that both measured and predicted AMPO were based on (near-)identical knee joint movements. Averaged across all participants, conditions and cycles, the sum of squared differences between the desired and measured knee joint angle was only (7 ± 12) · 10^-3^ rad. As a result, knee extension lasted only 0.6 ± 0.5 ms shorter than desired, and knee flexion lasted 0.6 ± 0.5 ms shorter than desired. In addition, the difference between maximum and minimum knee joint angle in the experiment was (0.7 ± 0.2) · 10^-3^ rad higher than desired, averaged across all participants, conditions and cycles. Overall, the dynamometer successfully imposed the desired knee joint movements both when participants were passive and when they aimed to maximise AMPO.

### 3.3 Timing of *m. quadriceps femoris* activation was adequate

EMG values during knee extension averaged over all conditions and participants were 78.1 ± 30.8% of *EMG^MVC^* for *m. quadriceps femoris* and 16.7 ± 12.2% of *EMG^MVC^* for the hamstring muscle group, between a knee joint angle of 0.45 and 1.55 rad. There was substantial interindividual variance in EMG values for both muscles, while variance in EMG values between conditions within each participant was small. EMG values of the hamstring muscle group increased just before the transition from flexion to extension and followed a similar time course as the EMG of *m. quadriceps femoris* (see Figure 5). This was reflected by a correlation coefficient of 0.77 ± 0.18 between EMG values of *m. quadriceps femoris* and of the hamstring muscle group, suggesting possible crosstalk between the two muscle groups. During flexion, EMG values of the hamstring muscle group were close to zero for all participants and conditions. EMG values of *m. quadriceps femoris* increased just before the transition from flexion to extension and were close to zero around the transition from extension to flexion for all participants and conditions (see Figure 5). Consequently, *M_mus_* rose around the transition from flexion to extension and was close to zero around the transition from extension to flexion (Figure 6). Except for occasional delays in the activation onset of *m. quadriceps femoris* during the first active cycle (cycle 6), *M_mus_* exhibited little variance between cycles (see also Figure 7). In sum, participants were capable of finding an activation of their *m. quadriceps femoris* that appeared to be (near-)optimal.

**Figure 5:**
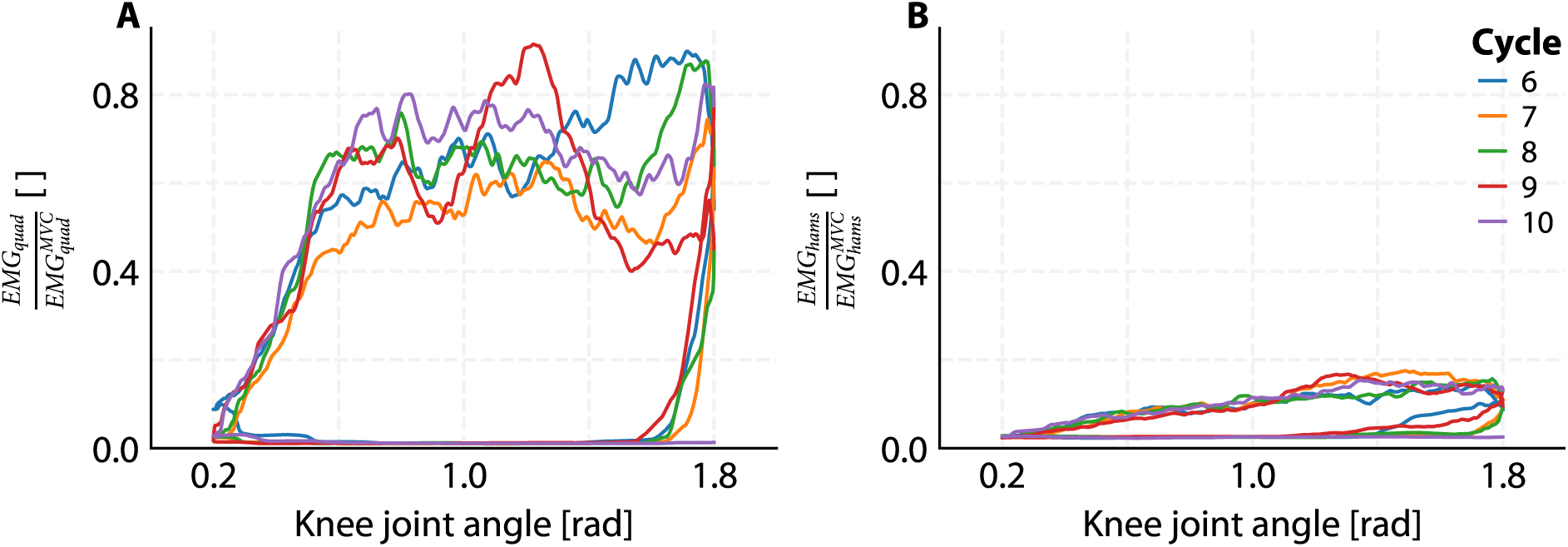
Representative example of measured EMG, shown for participant 5 under condition 0.35C. A) EMG of *m. vastus medialis* normalised by its *EMG^MVC^* value as a function of knee joint angle. EMG values increased just before the transition from flexion to extension and were close to zero during flexion, indicating that the timing of *m. quadriceps femoris* activation of participants was adequate. B) EMG of the *m. biceps femoris* normalised by its *EMG^MVC^* value as a function of knee joint angle. EMG values increased just before the transition from flexion to extension, following a similar time course as the EMG values of *m. vastus medialis*, suggesting possible co-contraction. During flexion, EMG values were close to zero, indicating participants adequately relaxed their the hamstring muscle group during flexion.

**Figure 6:**
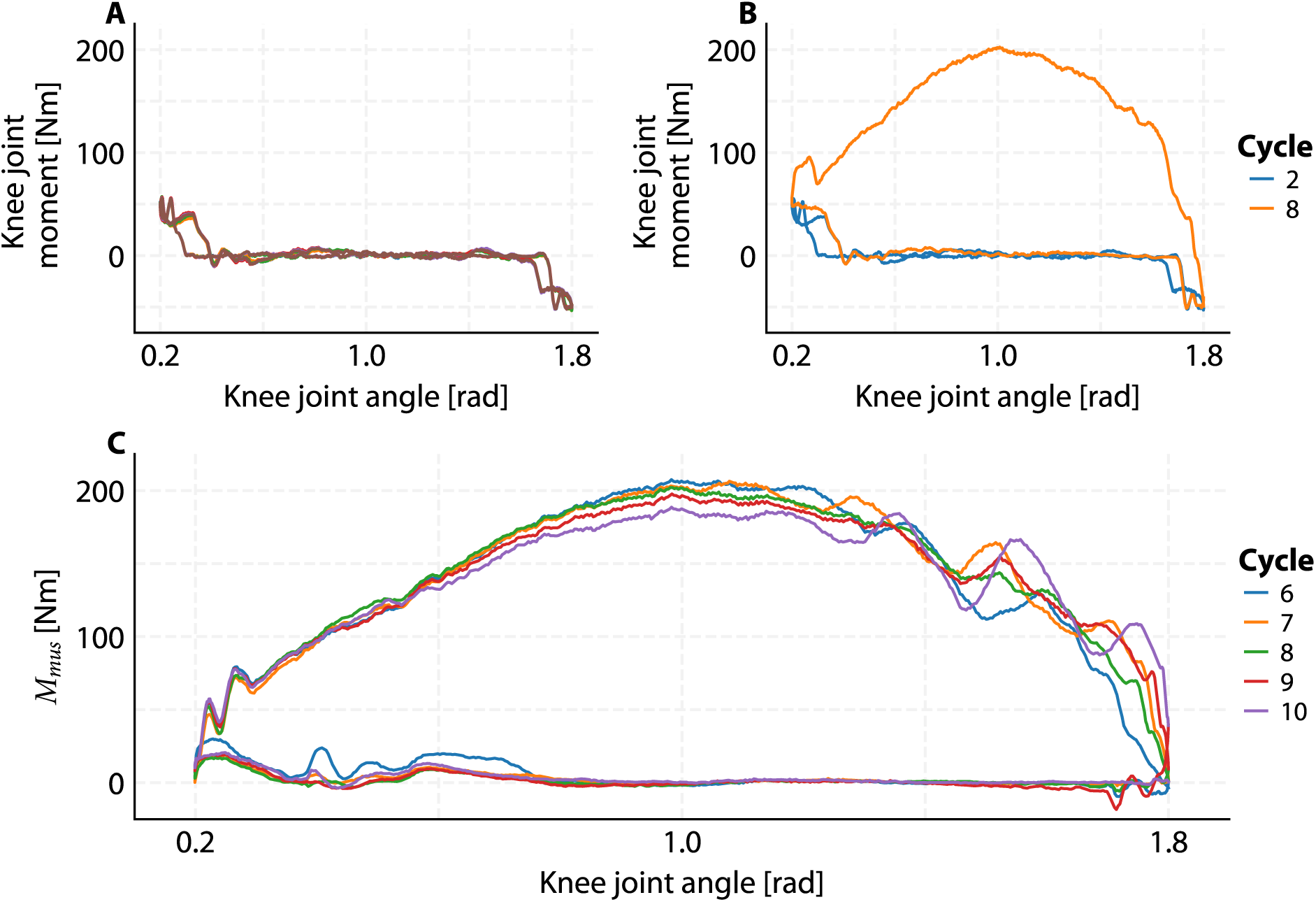
Representative example of mechanical behaviour, shown for participant 5 under condition 0.35C. A) Knee joint moment during passive cycles. Every individual cycle is represented by a different coloured line. B) Knee joint moment during (passive) cycle 2 (blue line) and during (active) cycle 8 (orange line). C) ‘Muscular’ knee joint moment (*M_mus_*) as a function of knee joint angle. *M_mus_* was calculated by subtracting average knee joint moment during the passive cycles from the knee joint moment during the active cycles (for details Section 2.3). Every individual cycle is represented by a different coloured line. *M_mus_* was close to zero during flexion, indicating that the net mechanical work was almost exclusively the result of mechanical work during extension.

**Figure 7:**
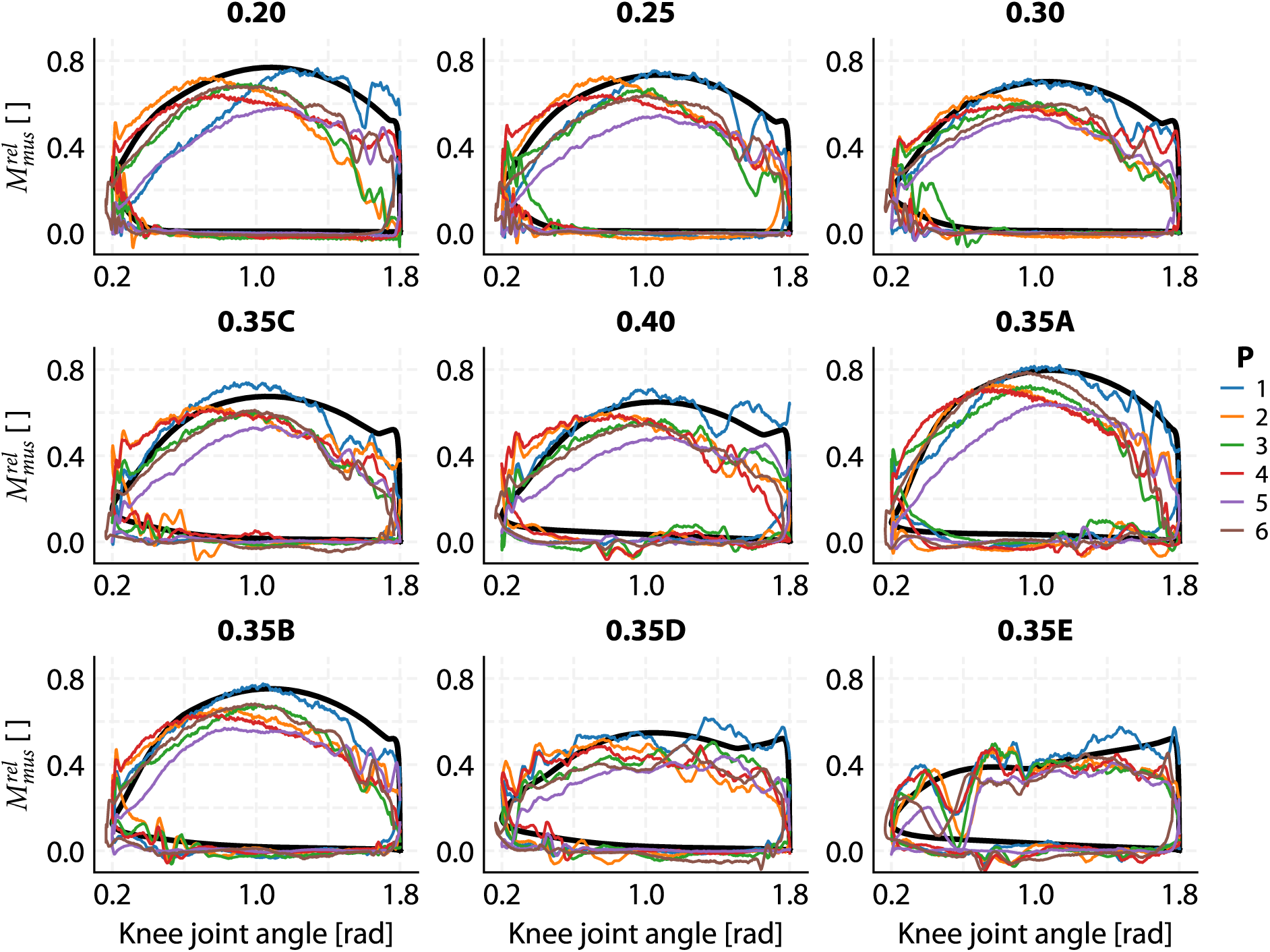
Experimentally estimated and model-predicted relative ‘muscular’ knee joint moment) for all conditions. *M^rel^* was calculated by dividing the ‘muscular’ knee joint moment by the maximal isometric knee joint moment of the individual participants (estimated from the isometric knee joint moment-angle relationship; see Figure 4) or by that of the model. The thick black line depicts the predicted *M^rel^*, while the thinner coloured lines depicts *M^rel^* for each individual participant (P). For each participant, *M^rel^* is shown for the cycle with highest measured AMPO.

### 3.4 Mechanical work was achieved almost exclusively during knee extension

Participants were instructed to maximise mechanical work per cycle using their *m. quadriceps femoris*, while keeping their hamstring muscle group completely relaxed at all times. Average net knee joint moment during the passive cycles was subtracted from net knee joint moment during the active cycles to correct for knee joint moment due to inertia, gravity and passive structures (see Figure 6). The resulting knee joint moment (*M_mus_*) was assumed to be predominantly the result of muscle force and was, as expected, substantially higher than zero during knee extension for all participants and conditions. Due to activation and deactivation delays, *M_mus_* may be above zero just before the transition from flexion to extension and immediately after the transition from extension to flexion, in order to maximise AMPO (see also Figure 7). Yet, *M_mus_* should be ideally zero during the middle part of knee flexion. For the active cycle the absolute value of *M_mus_* was 3.8 ± 4.7 Nm between a knee joint angle of 0.45 and 1.55 rad (corresponding to 100 ms after and before the transitions). This minimal *M_mus_* was a result of low activity of both *m. quadriceps femoris* (3.2 ± 3.3% of *EMG^MVC^*) and hamstring muscle group (4.6 ± 4.6% of *EMG^MVC^*) during the same time interval. We therefore concluded that participants effectively relaxed their muscles during knee flexion, such that the net mechanical work was almost exclusively the result of positive mechanical work delivered during knee extension (see Table 2).

**Table 2:**
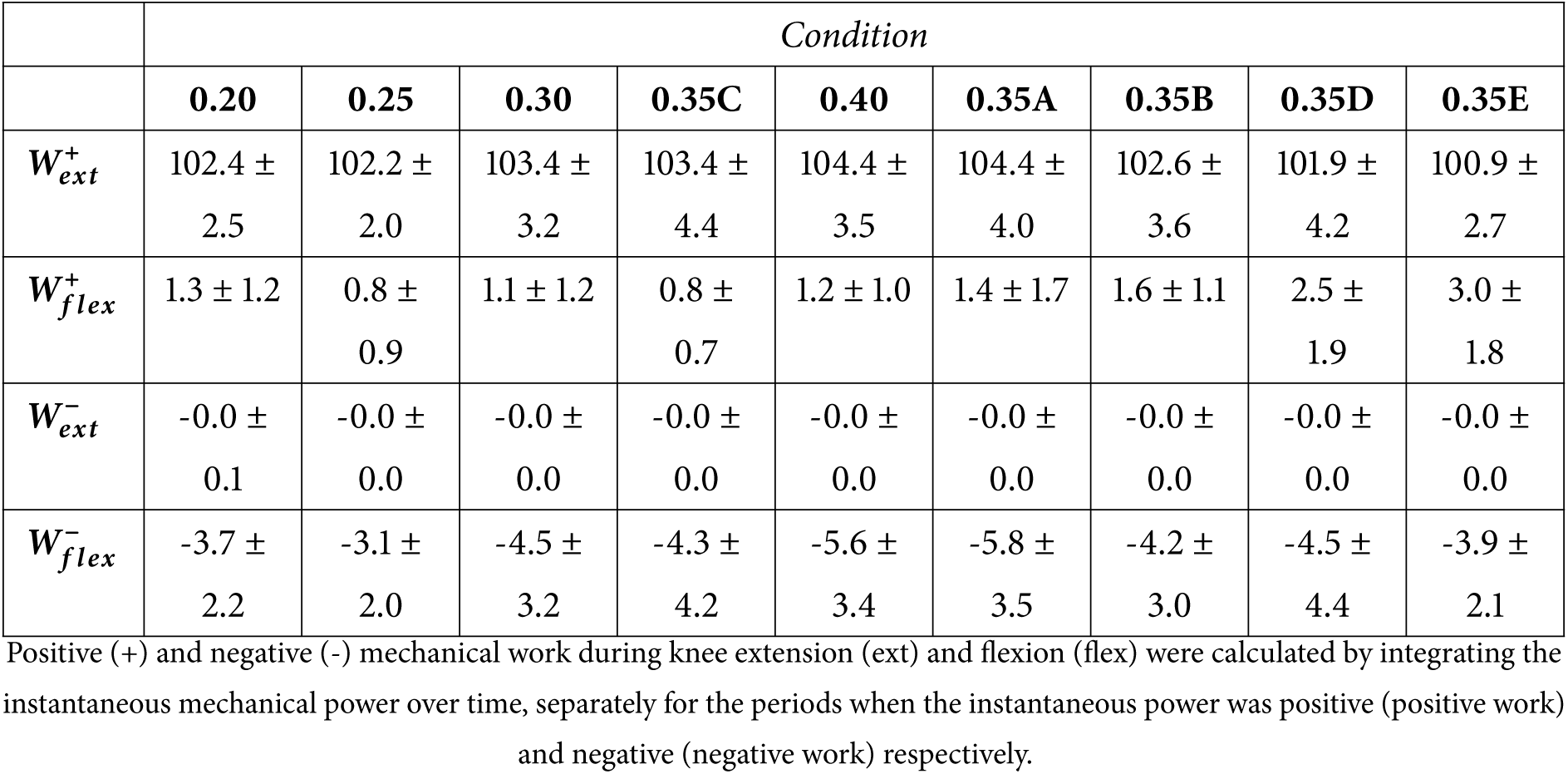
Positive and negative mechanical work during knee extension and flexion as a percentage of the net mechanical work per cycle.

### 3.5 Measured AMPO closely matched predicted maximally attainable AMPO

To assess the reliability of AMPO measurements, we examined the variability of measured AMPO across cycles and repeated conditions. Averaged across each condition and for each participant, the second-highest measured AMPO was only 2.4 ± 2.3% lower than the highest value, indicating low variance in AMPO between cycles. Furthermore, in repeated conditions, the highest measured AMPO at the end of the experiment was just 2.4 ± 4.3% lower than at the start. Together, these results indicate that AMPO measurements were consistent across cycles and throughout the experimental session.

Measured AMPO was compared against the predicted maximally attainable AMPO. First, AMPO was normalised by the generic value of *L^opt^_CE_* and participants’ individually estimated *F^max^_CE_* (see Section 2.3). As predicted, (1) measured AMPO differed substantially between conditions 0.20, 0.25, 0.30, 0.35 and 0.40 due to the imposed variations in cycle frequency and FTS and (2) measured AMPO was almost identical for conditions 0.35A-E, despite substantial variations in cycle frequency and FTS (Figure 8; Table 3). Overall, measured AMPO normalised by *L^opt^_CE_* and participants’ individual *F^max^_CE_* correlated strongly with predicted maximally attainable AMPO (r^2^=0.76). Second, AMPO was normalised by z-scores of measured and predicted maximally attainable AMPO. This revealed a very strong correlation between predictions and measurements (r^2^=0.95). All in all, measured AMPO was remarkably close to the predicted maximally attainable AMPO.

**Figure 8:**
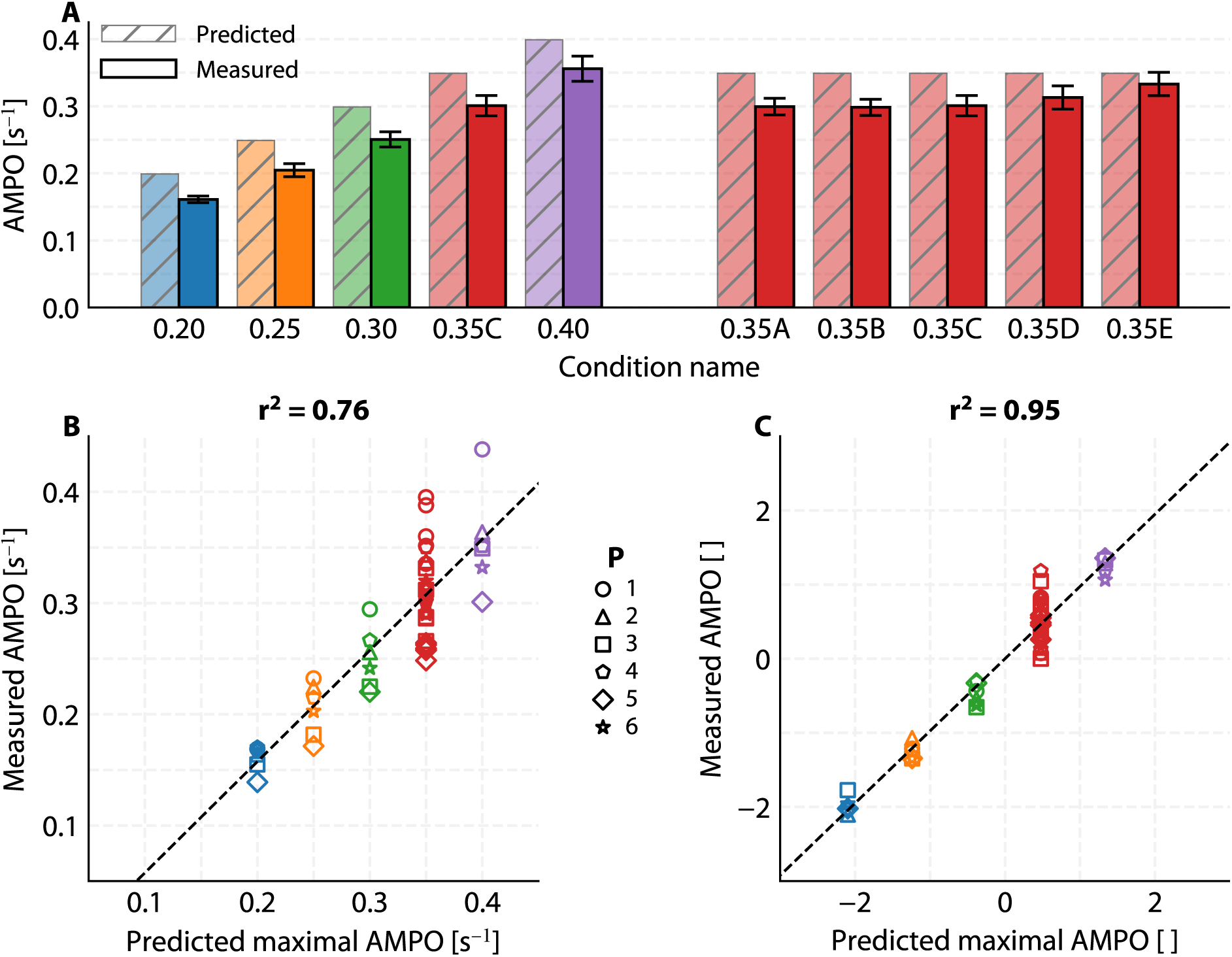
Measured AMPO versus predicted maximally attainable AMPO. A) Barplot showing predicted values with diagonal hatching and semi-transparent fill (*α* = 0.5), and experimentally measured values fully filled. Error bars denote the standard error of the mean. B) Scatterplot of measured versus predicted maximally attainable AMPO normalised by *L^opt^_CE_* and *F^max^_CE_*, with *F^max^_CE_* estimated individually for each participant. C) Scatterplot of measured versus predicted maximally attainable AMPO normalised to z-scores. Participants (P) are distinguished by symbols.

**Table 3:**
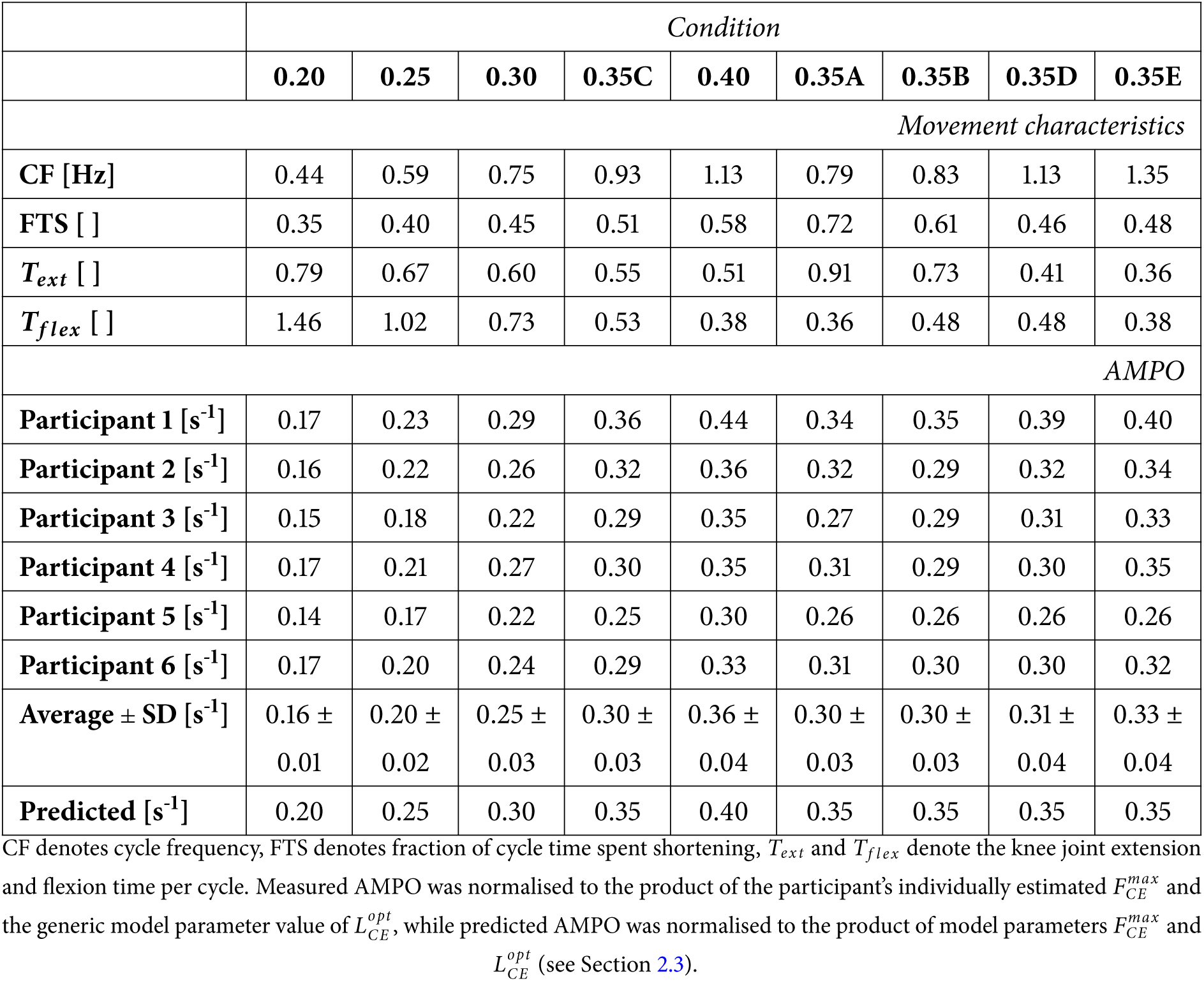
Overview of experimental conditions and measured AMPO per participant and condition.

|  | Condition |  |  |  |  |  |  |  |  |
| --- | --- | --- | --- | --- | --- | --- | --- | --- | --- |
|  | 0.20 | 0.25 | 0.30 | 0.35C | 0.40 | 0.35A | 0.35B | 0.35D | 0.35E |
| *Movement characteristics* |  |  |  |  |  |  |  |  |  |
| CF [Hz] | 0.44 | 0.59 | 0.75 | 0.93 | 1.13 | 0.79 | 0.83 | 1.13 | 1.35 |
| FTS [ ] | 0.35 | 0.40 | 0.45 | 0.51 | 0.58 | 0.72 | 0.61 | 0.46 | 0.48 |
| $T_{ext}$ [ ] | 0.79 | 0.67 | 0.60 | 0.55 | 0.51 | 0.91 | 0.73 | 0.41 | 0.36 |
| $T_{flex}$ [ ] | 1.46 | 1.02 | 0.73 | 0.53 | 0.38 | 0.36 | 0.48 | 0.48 | 0.38 |
| *AMPO* |  |  |  |  |  |  |  |  |  |
| Participant 1 [$s^{-1}$] | 0.17 | 0.23 | 0.29 | 0.36 | 0.44 | 0.34 | 0.35 | 0.39 | 0.40 |
| Participant 2 [$s^{-1}$] | 0.16 | 0.22 | 0.26 | 0.32 | 0.36 | 0.32 | 0.29 | 0.32 | 0.34 |
| Participant 3 [$s^{-1}$] | 0.15 | 0.18 | 0.22 | 0.29 | 0.35 | 0.27 | 0.29 | 0.31 | 0.33 |
| Participant 4 [$s^{-1}$] | 0.17 | 0.21 | 0.27 | 0.30 | 0.35 | 0.31 | 0.29 | 0.30 | 0.35 |
| Participant 5 [$s^{-1}$] | 0.14 | 0.17 | 0.22 | 0.25 | 0.30 | 0.26 | 0.26 | 0.26 | 0.26 |
| Participant 6 [$s^{-1}$] | 0.17 | 0.20 | 0.24 | 0.29 | 0.33 | 0.31 | 0.30 | 0.30 | 0.32 |
| Average ± SD [$s^{-1}$] | 0.16 ±<br>0.01 | 0.20 ±<br>0.02 | 0.25 ±<br>0.03 | 0.30 ±<br>0.03 | 0.36 ±<br>0.04 | 0.30 ±<br>0.03 | 0.30 ±<br>0.03 | 0.31 ±<br>0.04 | 0.33 ±<br>0.04 |
| Predicted [$s^{-1}$] | 0.20 | 0.25 | 0.30 | 0.35 | 0.40 | 0.35 | 0.35 | 0.35 | 0.35 |

CF denotes cycle frequency, FTS denotes fraction of cycle time spent shortening, $T_{ext}$ and $T_{flex}$ denote the knee joint extension and flexion time per cycle. Measured AMPO was normalised to the product of the participant's individually estimated $F_{CE}^{max}$ and the generic model parameter value of $L_{CE}^{opt}$ , while predicted AMPO was normalised to the product of model parameters $F_{CE}^{max}$ and $L_{CE}^{opt}$ (see Section 2.3).

### 3.6 Influence of SSC parameters on the maximally attainable AMPO

Having established that the influence of SSC parameters - cycle frequency, FTS and knee joint excursion - on the maximally attainable AMPO was accurately predicted by our Hill-type MTC model, we now examine the effects of the SSC parameters in detail. The predicted effect of cycle frequency and FTS at four distinct knee joint excursions are depicted in Figure 9. The optimal combination of SSC parameters was predicted to be a cycle frequency of about 1.6 Hz, a FTS of about 0.80 and a knee joint excursion of about 2.0 rad. Because knee joint movements were centred around 1.0 rad, knee joint excursions beyond 2.0 rad would result in knee hyperextension and were therefore not physiologically meaningful. Nevertheless, model predictions showed that knee joint excursions above 2.0 rad did not further increase the maximally attainable AMPO.

**Figure 9:**
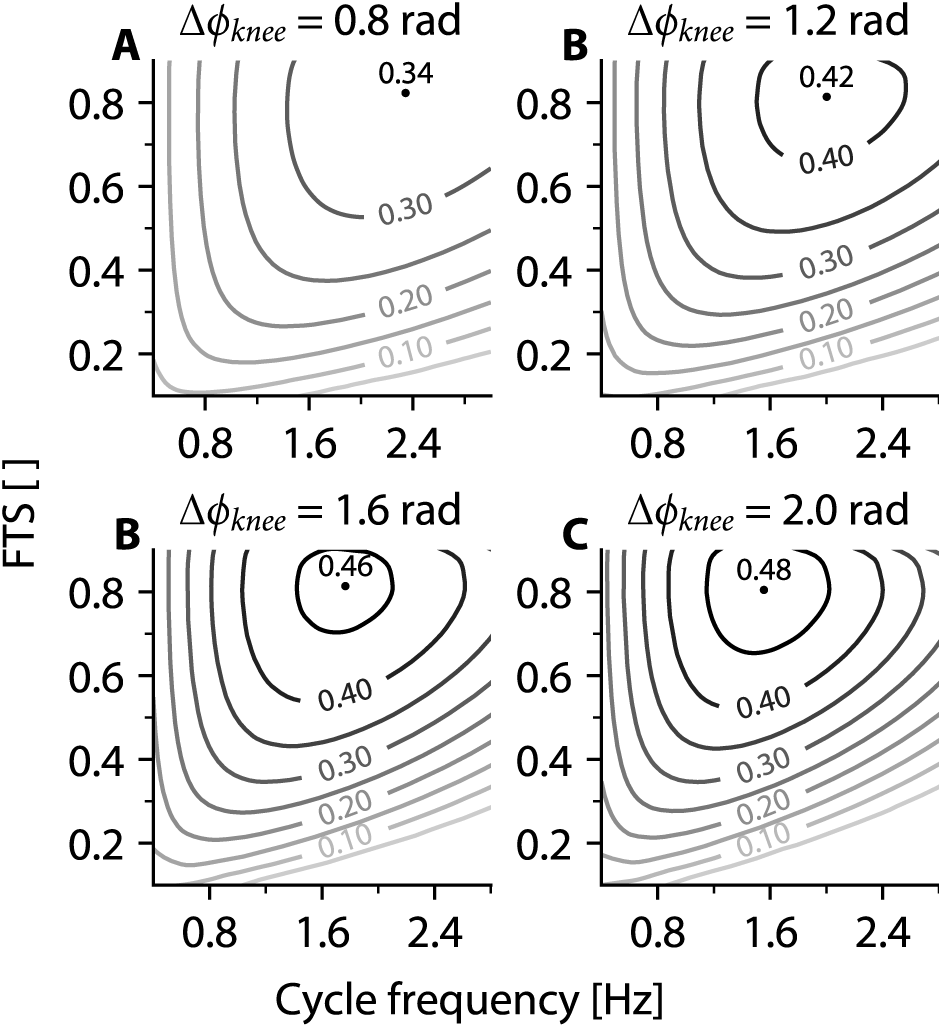
Predicted maximally attainable AMPO as a function of cycle frequency and fraction of the cycle time shortening (FTS) for four distinct knee joint excursions (Δ*ϕ_knee_*). Different values of the maximally attainable AMPO (normalised to *L^opt^_CE_* and *F^max^_CE_*) are depicted as individual contour lines. The dots indicate the location of peak AMPO corresponding to the optimal combination of cycle frequency and FTS for each knee joint excursion, with the corresponding peak AMPO labelled at the top-left of the dots.

Regarding the interrelationship between the SSC parameters, we found that the optimal FTS for AMPO remained relatively stable across the range of cycle frequencies and knee joint excursions studied, consis-tently ranging between 0.78 and 0.85. The largest variation in optimal FTS was observed at the smallest knee joint excursion studied (0.8 rad). By contrast, optimal cycle frequency showed substantial sensitivity to changes in FTS. Additionally, a strong interaction was found between cycle frequency and knee joint excursion: increasing one necessitates a decrease in the other to maximise AMPO.

## 4 Discussion

In this study, we examined the influence of imposed knee joint movement on the maximally attainable AMPO of human *m. quadriceps femoris* by combining experiments with simulations and optimisations using a Hill-type MTC model. Our findings revealed a strong correlation between measured AMPO and predicted maximally attainable AMPO, for a wide variety of imposed knee joint movements. Model predictions revealed that the maximally attainable AMPO peaks at a cycle frequency of about 1.6 Hz, an FTS of about 0.80, and a knee joint excursion of about 2.0 rad. While the optimal knee joint excursion was predicted to decrease with increasing cycle frequency and vice versa, the optimal FTS was predicted to be largely unaffected by changes in cycle frequency and knee joint excursion.

### 4.1 Evaluation of model predictions against experimental data

Measured AMPO and predicted maximally attainable AMPO were strongly correlated, with measured AMPO being 86 ± 10% of the predicted values on average. The lower measured AMPO primarily resulted from *M_mus_* values that were lower than values predicted for knee extension. It should be noted that, in theory, predicted maximally attainable AMPO should always exceed measured AMPO. For the model predictions, *m. quadriceps femoris* activation was set to be optimal in both timing and magnitude, and no hamstring co-contraction occurs. By contrast, participant muscle activation is inherently suboptimal.

Regarding *m. quadriceps femoris* activation, EMG of *m. quadriceps femoris* rose just at the transition from flexion to extension, causing *M_mus_* to increase at this time instant. Thus, we consider the timing of *m. quadriceps femoris* of participants to be adequate. However, EMG of *m. quadriceps femoris* did not exceed 100% of *EMG^MVC^* for all conditions amongst five of the six participants. This differs from previous studies (e.g. Bobbert and Harlaar, 1993; Kellis and Baltzopoulos, 1998; Westing et al., 1991), which reported *m. quadriceps femoris* EMG values slightly exceeding 100% in knee dynamometer experiments. Hence, submaximal *m. quadriceps femoris* activation may partly explain the lower *M_mus_* relative to predicted values.

Regarding the co-contraction of the hamstring muscle group, EMG of the hamstring muscle group during knee extension was 17.3 ± 12.4% of *EMG^MVC^*, consistent with previous studies (e.g. Kellis and Baltzopoulos, 1998; Snow et al., 1993). When interpreting surface EMG as a proxy for muscle activation, potential crosstalk should be considered. In the context of this study, it is possible that both quadriceps and hamstrings activation are slightly overestimated due to crosstalk. However, the effect of EMG crosstalk on the net knee joint moment is probably smaller than suggested by EMG values alone, since the maximal isometric force of the hamstring muscle group is approximately 2.5 times lower than that of *m. quadriceps femoris* (e.g. Prietto and Caiozzo, 1989). Nevertheless, it is clear that co-contraction of hamstrings during knee extension may also partly explain the slight deficit in *M_mus_* relative to predicted values.

Measured and predicted *M_mus_* were normalised to the maximal isometric force (*F^max^_CE_*; see Figure 7). *F^max^_CE_* was estimated individually for each participant from their measured isometric knee joint angle-moment relationship. In most cases, the actual participant’s maximum measured isometric knee joint moment was higher than the maximal value of the model (see Figure 4), indicating that *F^max^_CE_* was not overestimated. Nevertheless, it should be noted that any inaccuracy in *F^max^_CE_* directly affects normalised *M_mus_*. Aside from a simple normalisation of AMPO with *F^max^_CE_*, predictions were not individually tailored. Consequently, properties of participants’ *m. quadriceps femoris* could be different from the properties used in the Hill-type MTC model. Measured force rise was generally slower than predicted (see Figure 7), suggesting that the model’s activation dynamics may have been slightly too fast. Moreover, measured *M_mus_* was consistently lower across various knee joint angles, even where maximal activation was achieved in both experiment and model. The force–length relationship likely played only a minor role, because the isometric knee joint moment–angle relationship was reasonably similar for participants and the model. Instead, the force–velocity relationship appears to be the main contributor. In our model, the decline in knee extension moment with increasing angular velocity was at the lower end of experimentally observed values (for review, see Chen and Franklin, 2025). Overall, the slight deficit in *M_mus_* relative to predicted values may partly be explained by differences between the generic model properties and participants’ *m. quadriceps femoris* properties. We expect that, with individually tailored models, differences between experimental and predicted *M_mus_* would have been even smaller than differences reported in this study.

We emphasise that our aim was not to precisely predict the magnitude of *M_mus_* over time, but rather to examine the influence of SSC parameters on the maximally attainable AMPO. Thus, it is promising that we predict the magnitude of the maximally attainable AMPO remarkably well using a relatively simple Hill-type MTC model with generic parameter values obtained from literature, despite the aforementioned methodological limitations. Moreover, when assessing the influence of SSC parameters on the maximally attainable AMPO, the relative change in AMPO is more informative than its (average) magnitude. In this context, it is particularly encouraging that AMPO normalised to z-scores revealed a very strong correlation between predictions and measurements (r^2^ = 0.95). This finding aligns with previous studies demonstrating that Hill-type MTC models reliably predict mechanical behaviour, especially during concentric contractions (e.g. Ingen Schenau et al., 1988; Lemaire et al., 2016; Sandercock and Heckman, 1997; Wakeling and Johnston, 1999; Zandwijk et al., 1996). Concentric contractions primarily determine maximally attainable AMPO. All in all, we attribute the small discrepancy between measured and predicted maximally attainable AMPO to (a combination of) several factors, such as suboptimal *m. quadriceps femoris* activation, hamstring co-contraction, and generic model assumptions. Nevertheless, predictions aligned remarkably well with measurements, demonstrating the validity of the predicted influence of SSC parameters on the maximally attainable AMPO of human *m. quadriceps femoris*.

### 4.2 Results agree well with available literature

In the first part of the current study, *L*^0^_SEE_, *L^opt^_CE_* and *F^max^_CE_* were estimated for each participant individually based on isometric measurements at nine different knee joint angles. The measured net knee joint moment-angle relationship of participants was in line with literature (e.g. Anderson et al., 2007; Eijden et al., 2008; Murray et al., 1980). The estimation of the muscle model parameters was deemed successful as the root mean square error between the measured isometric knee joint moment and the musculoskeletal predicted isometric knee joint moment was low (6.6 ± 2.2% of *F^max^_CE_* on average). Additionally, values of *L*^0^_SEE_ and *L^opt^_CE_*corresponded well with values obtained from cadavers (e.g. Friederich and Brand, 1990; Ward et al., 2009; Wickiewicz et al., 1983) and values used in other modelling studies (e.g. Anderson and Pandy, 1999; Soest and Bobbert, 1993). Overall, we conclude that the values of *L*^0^_SEE_, *L^opt^_CE_* and *F^max^_CE_* were successfully estimated.

In the second part of the current study, we examined the maximally attainable AMPO for a large set of imposed knee joint movements. As our study is the first to investigate this for human muscle, the only available data for comparison are those of *in vitro* or *in situ* studies on animal muscle. Whereas we influenced muscle length indirectly by imposing knee joint excursions, those studies directly imposed muscle length excursions. Those studies on animal muscle show that an increase in cycle frequency leads to a decrease in the AMPO-optimal length excursion, muscle stimulation duration and mechanical work per cycle (e.g. Askew and Marsh, 1997; James et al., 1995; Josephson, 1985; Reuvers et al., 2025). Our findings for human *m. quadriceps femoris* align with these observations. Additionally, in accordance with Askew and Marsh (1997) and Reuvers et al. (2025), we found that the optimal (length) excursion increases with increasing FTS and decreasing cycle frequency.

Regarding the values of cycle frequency, FTS and muscle fibre length excursion per se, it should be noted that these values substantially differ due to differences in muscle properties (e.g. maximal shortening velocity, excitation dynamics etc.). Accordingly, the optimal cycle frequency of about 1.6 Hz observed for human *m. quadriceps femoris* contrasts with about 4 Hz for mouse *m. soleus* (Askew and Marsh, 1997), about 9 Hz for mouse *m. extensor digitorum longus* (Askew and Marsh, 1997), and about 3.5 Hz for rat *m. gastrocnemius medialis* (Reuvers et al., 2025). Similarly, optimal muscle fibre length excursion, expressed as a percentage of the optimum fibre length, varies across muscles: 22% for mouse *m. soleus* (Askew and Marsh, 1997), 16% for mouse *m. extensor digitorum longus* (Askew and Marsh, 1997), and 58% for rat *m. gastrocnemius medialis* (Reuvers et al., 2025). All of these values are considerably lower than the 85% reported here for human *m. quadriceps femoris*. Interestingly, the optimal FTS of 0.80 for human *m. quadriceps femoris* is comparable to the range of 0.80-0.90 observed in animal muscle (Askew and Marsh, 1997; Reuvers et al., 2025). The optimal FTS is predominantly a trade-off between (the negative effect of) the force-velocity relationship on the one hand and (the negative effect of) the excitation dynamics on the other hand (for details, see Reuvers et al., 2025). Both factors depend on muscle fibre type; their negative effect on AMPO increases with slower muscle fibre types. Consequently, the optimal FTS is expected to be relatively robust for variations in muscle fibre type. Therefore, it is promising that the optimal FTS in our study aligns with those observed in animal muscle. Overall, our findings are consistent with the available literature, which strengthens our confidence in the validity of the results.

### 4.3 Experimental constraints and physiological relevance

In extensive pilot experiments, we explored a range of knee joint movements with different angular velocities and accelerations. Based on these pilot experiments and safety considerations, we limited knee joint acceleration to 50 rad/s^2^, thereby also limiting the (maximum achievable) knee joint angular velocity. As a result, only a subset of combinations of SSC parameters — cycle frequency, FTS, and knee joint excursion — could be tested experimentally. Importantly, this does not imply that higher velocities or accelerations are physiologically infeasible for human knee joint movements. In fact, substantially higher values are reached during various motor activities (for review, see Grimmer et al., 2020). For example, during sprint cycling, knee joint angular velocities reach 5–10 rad/s, and angular accelerations exceed 100 rad/s^2^. During sprint running these values are even higher, with knee joint angular velocities reaching about 20 rad/s and angular accelerations up to 200 rad/s^2^. Therefore, the joint movements that were not feasible in our experiment are nonetheless physiologically meaningful.

### 4.4 Physiological implications for human movement

AMPO in our study was normalised to the product of *F^max^_CE_* and *L^opt^_CE_*. This allowed comparison across participants, but also allows us to extend our findings to muscles of similar fibre types. For typical values of *L^opt^_CE_*=9.3 cm and *F^max^_CE_* =5250 N for m. vastii, and *L^opt^_CE_*=8.1 cm and *F^max^_CE_* =1750 N for m. rectus femoris (Soest and Casius, 2000), peak AMPO for *m. quadriceps femoris* is predicted to be approximately 300 W per leg at the optimal combination of SSC parameters. This value may be used as a benchmark in assessing maximum AMPO of human *m. quadriceps femoris* AMPO in any short-duration task.

Figure 9 shows that SSC parameters have substantial influence on the maximally attainable AMPO across a realistic range of knee joint movement. For example, decreasing FTS from its optimal value of about 0.8 to 0.5 reduces AMPO by about 20% at the optimal cycle frequency, with smaller effects at lower frequencies and larger effects at higher frequencies. In many periodic sports, *m. quadriceps femoris* shortening duration does not exceed the lengthening duration. For example, FTS ranges from 0.3-0.5 in rowing (Bechard et al., 2009; Ettema et al., 2022), is about 0.3 in cross-country skiing (Nilsson et al., 2004; Pellegrini et al., 2014), and is about 0.5 in cycling (Bobbert et al., 2020). These suboptimal FTS values are often due to equipment design, highlighting the potential performance gains from designs that allow muscles to operate closer to their optimal SSC parameters.

### 4.5 Conclusion

In this study, we examined how SSC parameters — cycle frequency, FTS, and knee joint excursion — influence the maximally attainable AMPO of human *m. quadriceps femoris* during short-duration, all-out (sprint) conditions. Because only a subset of SSC parameters can be tested experimentally, we combined experiments with musculoskeletal modelling. Differences in maximally attainable AMPO were adequately predicted by our Hill-type MTC model, justifying its use to examine the effect of SSC parameters on maximally attainable AMPO in detail. Model predictions showed a substantial influence of SSC parameters on the maximally attainable AMPO. Specifically, predictions revealed a strong interaction between cycle frequency and knee joint excursion: increasing one necessitates a decrease in the other to maximise AMPO. Even more interestingly, in order to maximise AMPO, *m. quadriceps femoris* should spend about 80% of the cycle duration while shortening, independent of cycle frequency and/or knee joint excursion.

## Acknowledgments

The authors thank all participants for their sustained and dedicated effort during the experiment.

## Data and resource availability

All data, code, and materials used in this study are openly available:

- **GitHub repository:** All raw data, processed data, and analysis code are hosted on GitHub at https://github.com/edwinreuvers/hq-ampo.
- **Reproducible analysis website:** Full analysis pipeline — including data, analysis code, and figure/table generation — is available at https://edwinreuvers.github.io/publications/hq-ampo.

## Appendix

### A1 Selection of experimental conditions

With our musculoskeletal model we predicted the maximally attainable AMPO for knee joint movements with different constant knee angular velocities during flexion and extension. Thus, the knee joint angular velocity changed instantaneously at the switch from flexion to extension (and vice versa), and the knee joint angular acceleration at that instant was infinitely high/low. Clearly, infinite knee joint accelerations cannot be achieved in an experimental setup. Instead, we chose to impose a constant knee angular acceleration until a predefined knee joint angular velocity was achieved — determined by the combination of cycle frequency, FTS, knee joint angle excursion and constant knee angular acceleration — in order to achieve the highest cycle frequency and the greatest difference between maximal and minimal FTS. In extensive pilot tests, we determined that 50 rad/s^2^ was the highest knee angular acceleration that participants were comfortable with and was used as an upper limit for the experiment. Accordingly, the experimental conditions were selected from the knee joint movement that were experimentally feasible (see Figure 3).

**Figure A1:**
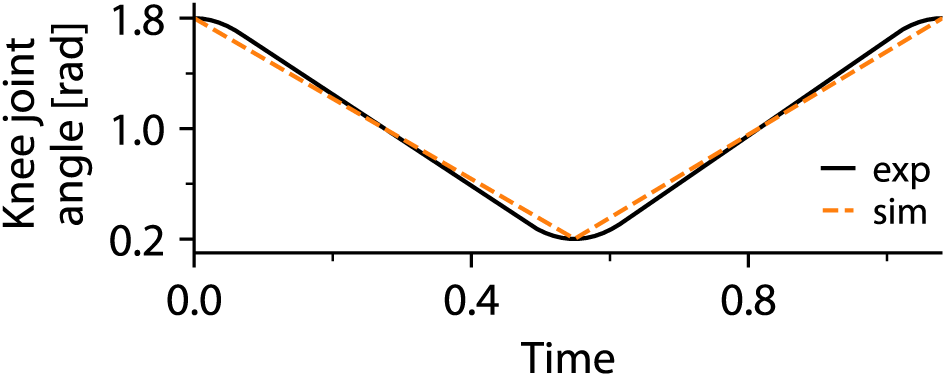
Comparison between knee joint movements during the experiment and simulations. In the experiment, knee joint movements were limited to a angular acceleration of 50 rad/s^2^. In contrast, in the simulations, knee joint movements were used with different constant angular velocities for flexion and extension in order to simulate a broad range knee joint movements.

Due to the limit on maximal knee joint acceleration, the imposed knee joint movement in the experiment were slightly different from those with a constant knee angular velocity (i.e. with no limit on knee angular acceleration; see Figure A1). As a result, the maximally attainable AMPO differed slightly between those two for the same set of SSC parameters. Therefore, we derived predictions of the maximally attainable AMPO for knee joint movement with a limit of 50 rad/ŝ2 on the knee angular acceleration (see Figure 3).

Among the experimentally feasible knee joint movements, we observed that the influence of cycle frequency and FTS was greatest at highest knee joint excursion. Therefore, we chose to use a knee joint excursion of 1.6 rad, which was the maximum range of motion of the knee dynamometer. Based on the resulting mapping from cycle frequency and FTS at a knee joint excursion of 1.6 rad (see contour lines in Figure 3), we selected two sets of five experimental conditions. One condition was shared between both sets, resulting in nine unique experimental conditions in total. The first set of five conditions was chosen such that the maximally attainable AMPO was predicted to be identical, despite substantial variations in cycle frequency and FTS (0.35A-E; Figure 3). The second set of conditions was chosen such that the maximally attainable AMPO was predicted to differ substantially due to the imposed variations in cycle frequency and FTS (0.20, 0.25, 0.30, 0.35A-E and 0.40; Figure 3).

### A2 Musculoskeletal model

In both our predictions, as well as in the experiments, the knee joint angle over time was fully imposed. Based on the imposed knee joint angle, we calculated the muscle–tendon complex (MTC) length over time using a quadratic relation, as first suggested by Grieve et al. (1978):

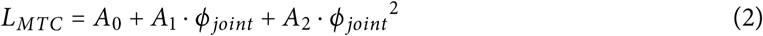

*A*_0_ denotes the muscle-tendon-complex length at a knee joint angle of 0 rad (full extension), with knee joint angle increasing with flexion. *ϕ_joint_* denotes the knee joint angle. Both *A*_1_ and *A*_2_ define the change in MTC length based on knee joint angle. MTC mechanical work can be calculated both by *F_MTC_dL_MTC_*and *M_MTC_dϕ_joint_* = *F_MTC_ r_MTC_dϕ_joint_*. An expression for the moment arm of the muscle-tendon-complex around the knee joint (*r_MTC_*) can then be derived by substituting these equations into Equation 2:

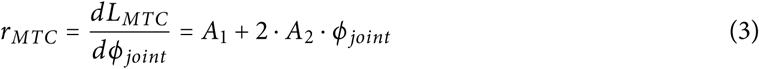

**Table A1:** Key muscle parameter values.

| Description | Symbol | Unit | Value | Reference |
| --- | --- | --- | --- | --- |
| *Musculoskeletal geometry* |  |  |  |  |
| MTC length at $\phi_{knee} = 0$ | $A_0$ | m | 0.21 | Soest and Casius (2000) |
| Linear coefficient of MTC-$\phi_{knee}$ relation | $A_1$ | m/rad | 0.042 | Soest and Casius (2000) |
| Quadratic coefficient of MTC-$\phi_{knee}$ relation | $A_2$ | m/rad<sup>2</sup> | 0 | Soest and Casius (2000) |
| *Contraction dynamics* |  |  |  |  |
| Hill constant | $a^{rel}$ | - | 0.41 | Soest and Casius (2000) |
| Hill constant | $b^{rel}$ | 1/s | 5.2 | Soest and Casius (2000) |
| Maximum isometric CE force | $F_{CE}^{max}$ | N | 5250 | Soest and Casius (2000) |
| SEE stiffness scaling parameter | $k_{SEE}$ | N/m<sup>2</sup> | $1.28 \cdot 10^8$ | Soest and Casius (2000) |
| CE optimum length | $L_{CE}^{opt}$ | m | 0.093 | Soest and Casius (2000) |
| SEE slack length | $L_{SEE}^0$ | m | 0.16 | Soest and Casius (2000) |
| Determines CE force-length relation width | $w$ | - | 0.56 | Soest and Casius (2000) |
| *Excitation dynamics* |  |  |  |  |
| Determines steepness γ-$q$ relation | $a_{act}$ | - | -4.59 | Hatze (1981) |
| Determines $pCa^{2+}$ level at which $q = 0.5$ | $b_{act,1}$ | - | 5.17 | Hatze (1981) |
| Determines $pCa^{2+}$ level at which $q = 0.5$ | $b_{act,2}$ | - | 1.08 | Hatze (1981) |
| Determines $pCa^{2+}$ level at which $q = 0.5$ | $b_{act,3}$ | - | -0.19 | Hatze (1981) |
| Relates $\gamma$ to $pCa^{2+}$ | $kCa$ | mol/L | $8.00 \cdot 10^{-6}$ | Kistemaker et al. (2005) |
| Activation time constant | $\tau_{act}$ | s | $88.9 \cdot 10^{-3}$ | Hatze (1981) |
| De-activation time constant | $\tau_{deact}$ | s | $88.9 \cdot 10^{-3}$ | Hatze (1981) |

Parameter values are depicted only for those key to predict the maximally attainable AMPO.

